# Joint Registration of Multiple Point Clouds for Fast Particle Fusion in Localization Microscopy

**DOI:** 10.1101/2021.09.09.459453

**Authors:** Wenxiu Wang, Hamidreza Heydarian, Teun A.P.M. Huijben, Sjoerd Stallinga, Bernd Rieger

## Abstract

We present a fast particle fusion method for particles imaged with single-molecule localization microscopy. The state-of-the-art approach based on all-to-all registration has proven to work well but its computational cost scales unfavourably with the number of particles *N*, namely as *N*^2^. Our method overcomes this problem and achieves a linear scaling of computational cost with *N* by making use of the Joint Registration of Multiple Point Clouds (JRMPC) method. Straightforward application of JRMPC fails as mostly locally optimal solutions are found. These usually contain several overlapping clusters, that each consist of well-aligned particles, but that have different poses. We solve this issue by repeated runs of JRMPC for different initial conditions, followed by a classification step to identify the clusters, and a connection step to link the different clusters obtained for different initializations. In this way a single well-aligned structure is obtained containing the majority of the particles.

We achieve reconstructions of experimental DNA-origami datasets consisting of close to 400 particles within only 10 min on a CPU, with an image resolution of 3.2 nm. In addition, we show artifact-free reconstructions of symmetric structures without making any use of the symmetry. We also demonstrate that the method works well for poor data with a low density of labelling and for 3D data.

## 1 Introduction

The diffraction of light limits the resolution of conventional microscopy to about 200 nm. Several super-resolution microscopy techniques enable “diffraction unlimited” resolution [8, 16, 26]. Single-molecule localization microscopy (SMLM) is a widely used member of the family of super-resolution techniques, and obtains super-resolved images by localizing single fluorescent emitters. The resolution of these super-resolved images is not infinite, but in practice restricted to about 20 nm due to the incomplete fluorescent labelling and a limited number of collected photons per localization event [21]. In recent years, significant improvements have been made to increase the photon count per localization [20]. Increasing the density of labelling (*DOL*) using biochemical means is difficult, where *DOL* values of around 50% are typically achieved. In addition, a high local *DOL* can lead to an increased rate of mislocalizations [5] which is detrimental for the quality of the imaging process. If the sample includes many chemically identical bio-complexes (called particles in the following), the limitation imposed by a low *DOL* can be lifted by fusion of all these particle into one single reconstruction, the so-called super-particle, leading to a much better resolution and signal-to-noise ratio (SNR) [18, 23]. This approach by particle fusion, of course, ignores potential heterogeneity in the underlying biology within the collection of particles. Template-driven particle fusion methods have been used [2, 7, 18, 23], but have a substantial risk of resulting in a biased reconstructed structure. Heydarian *et al*. proposed a template-free particle fusion method based on an all-to-all registration (all-to-all method in short), which is robust against underlabelling and misregistration [10, 13]. The all-to-all method has proven to work well and produces reconstruction resolutions down to a few nanometers. Despite this success, computational times of around a day for a number of particles *N* exceeding about 1000 − are not uncommon and are only feasible with the use of GPU acceleration. The root cause lies within the unfavourable scaling of computational cost with *N*^2^, because each particle is registered to all other particles, resulting in *N* (*N* − 1)/2 registration pairs. The all-to-all method has another drawback, the so-called “hot-spot” problem. For symmetric structures, random variations in the localization data with binding site are amplified by the pair-wise optimal registration process. Heydarian *et al*. solved this problem by first detecting the present symmetry and then imposing it on the data in a post-processing step. Thus, a particle fusion algorithm that is fast and which avoids the hot-spot artifact is desired.

An alternative to the all-to-all method is based on the Joint Registration of Multiple Point Clouds (JRMPC) method [4]. In the JRMPC method, particles are iteratively rotated and translated to fit to a Gaussian Mixtures Model (GMM), which is updated itself in each iteration round. The key advantage of the JRMPC method is that the computational complexity scales linearly with the number of particles *N*, which makes it inherently faster than the all-to-all method if *N* grows large. In addition, hot-spot artifacts in symmetric structures are avoided without imposing (a-priori) symmetry information, because the joint registration treats each particle equally. There are, however, major drawbacks to the JRMPC method. First, the outcome of the JRMPC turns out to be highly susceptible to the initialization of the GMM (number of Gaussians, center positions and widths). Different initial settings of the GMM parameters lead to different sets of final estimated particle rotations and translations. Second, the final outcome usually consists of several clusters, where the particles within the clusters are well-registered, but where the clusters have different poses. We attribute these issues with robustness of the algorithm to trapping in local optima of the iterative optimization (outlined in section 2.1 in detail).

The goal of the work presented in this paper is to overcome the robustness problems of the JRMPC method while maintaining the inherent speed advantage. To this end, we propose a processing pipeline in which we combine JRMPC registration outcomes obtained with different GMM initializations using cluster analysis tools. The cluster analysis uses our recent unsupervised classification framework [14], which is based on the Bhattacharya distance metric [2] together with multi-dimensional scaling (MDS) [19] and k-means clustering [3, 15]. The process of JRMPC and classification is repeated several times for different GMM initializations. Pairs of clusters from different initializations may share particles. The relative poses of such particles in different clusters is used in a final step to combine the different clusters into a single well-aligned structure.

## 2 Method

Our proposed algorithm has three main steps, illustrated in Figure 1. The steps are (1) alignment of particles with JRMPC using multiple initializations, (2) classification of JRMPC registered particles into clusters, and (3) connection of the identified clusters into a single final reconstruction.

**Figure 1.**
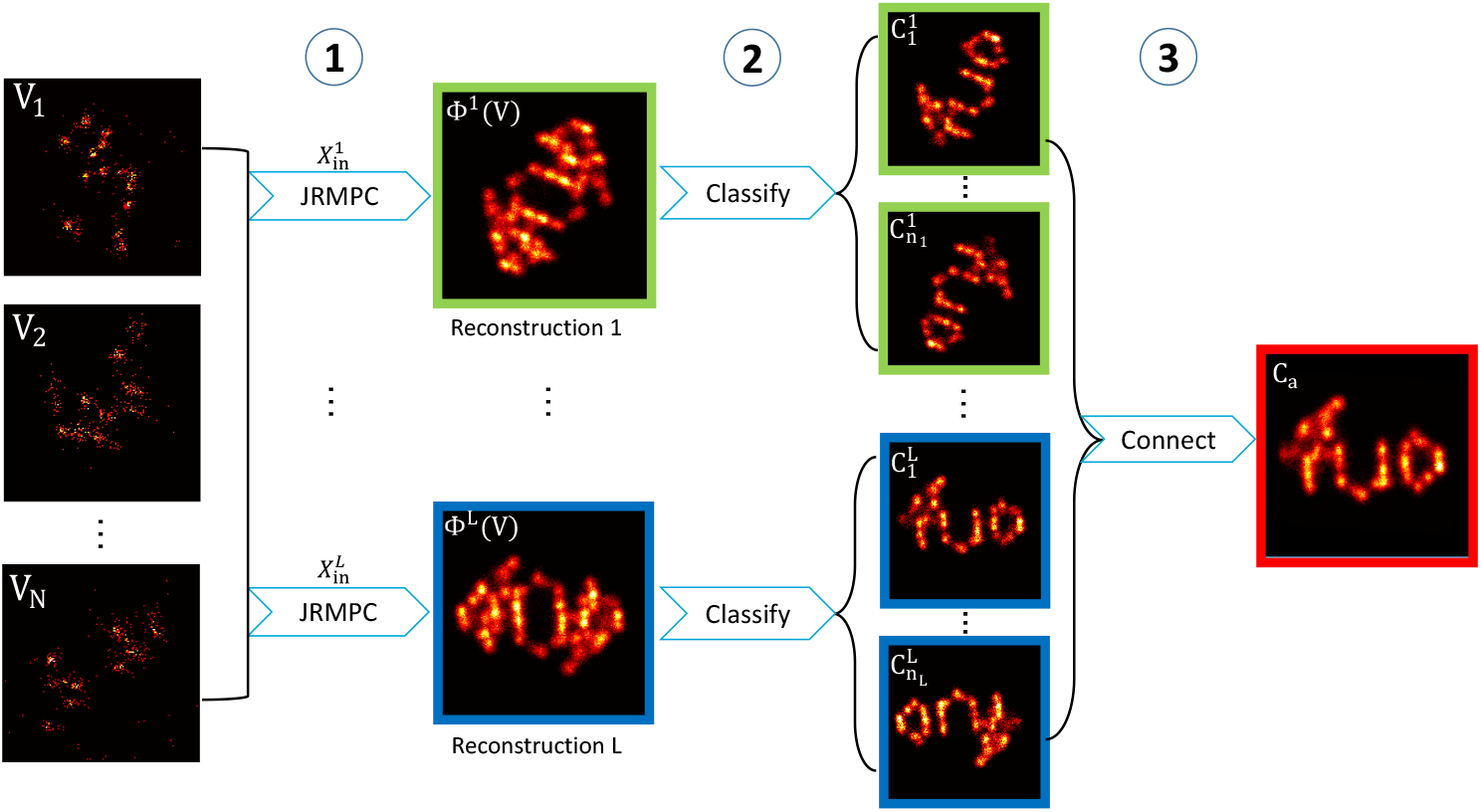
The three main steps of the proposed particle fusion algorithm. Step 1: Use JRMPC [4] to initially align *N* input particles 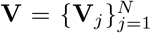 with *L* random initializations of the GMM 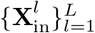 leading to *L* different reconstructions 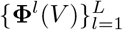. Step 2: Apply the unsupervised classification method of Huijben *et al*.[14] to classify each reconstruction Φ^*l*^(*V*) into *n*_*l*_ clusters 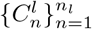 separating different overlapping poses in the reconstructed particles. Step 3: Connect particles from different clusters into the final super-particle reconstruction *C*_*a*_, such that each input particle is present at most once.

The input data is a union of particles 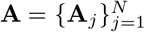, with *N* the number of particles. Each particle is characterized by a set of localization coordinates **V**_*j*_ and attendant localization uncertainties **Δ**_*j*_ as **A**_*j*_ = {**V**_*j*_; **Δ**_*j*_}. The coordinates of particle *j* represent *M*_*j*_ localizations:

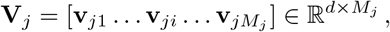

where the **v**_*ji*_ are vectors with elements equal to the *d* coordinates of the *i*-th localization in particle *j*. Depending on the data, the dimensionality *d* can be 2 or 3. In general, the localization uncertainties of the *M*_*j*_ localization events in particle *j* are:

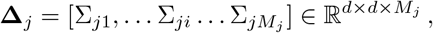

where the Σ_*ji*_ are *d* × *d* matrices equal to the covariance matrices of the *i*-th localization in particle *j*. Often a more simple description of the localization uncertainty is possible. For 2D data for example, the uncertainties are isotropic, and **Δ**_*j*_ can be written as:

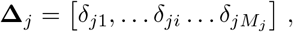

where the *δ*_*ji*_ are now scalar values that represent the localization uncertainty in the *xy* plane for the *i*-th localization in particle *j*. For most 3D data, **Δ**_*j*_ is represented as:

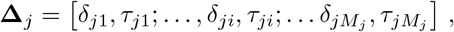

where now *τ*_*ji*_ is the localization uncertainty along the *z*-axis for the *i*-th localization in particle *j*. This axial localization uncertainty is typically larger than the uncertainty in the *xy* plane [22].

### 2.1 Alignment

The structure of the reconstruction is characterized in the JRMPC method by a GMM with parameters 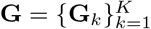, where each of the *K* Gaussians components **G**_*k*_ = [*p*_*k*_, ***µ***_*k*_, *σ*_*k*_] has a mixing coefficient (weight) *p*_*k*_, a set of *d* coordinates ***µ***_*k*_ that represent the mean of the Gaussian, and a standard deviation *σ*_*k*_ (an isotropic covariance matrix 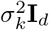 is taken). The GMM parameters have an initial setting **G**_in_, described in section 3.2. The parameters that are updated during the iterative JRMPC algorithm are:

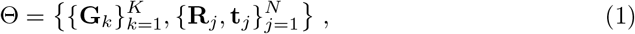

where *R*_*j*_ ∈ ℝ^*d×d*^ is the rotation applied to particle *j* and where *t*_*j*_ ∈ ℝ^*d×*1^ is the translation applied to particle *j*. The coordinates of the reconstruction are then:

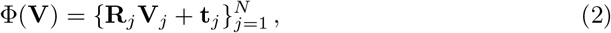

which thus contains the coordinates of all localization events in all particles. It is noted that the localization uncertainties are not taken into account in the JRMPC method. Further details on the steps in each iteration round of the JRMPC are given in Appendix A.

The outcome of the JRMPC depends on the choice of the initial GMM centers in **G**_in_. Our algorithm uses *L* differently initialized GMMs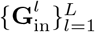, leading to *L* different JRMPC alignments 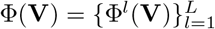 of the same union of particles with coordinates **V**.

### 2.2 Classification

The JRMPC algorithm can end up in a local optimum, resulting in multiple groups of particles (clusters) with different overlapping poses in the reconstruction. To separate these clusters, we use an unsupervised classification method recently proposed by our group [14]. This method enables the analysis of structural heterogeneity in localization datasets arising from e.g. naturally occurring biological variations. Here, we use this pipeline to decompose the *L* different JRMPC outcomes into clusters of particles, where the particles within each cluster are well-aligned. First, we compute the normalized Bhattacharya cost function between every transformed particle 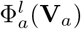 and every other transformed particle 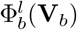 within the JRMPC registration for each initialization *l* = 1, 2, …, *L*. This one time computation gives an upper triangular matrix with *N* (*N* − 1)/2 cost function values *S*. The normalized Bhattacharya cost is in general given by the sum over the *M*_*a*_ localizations of particle *a* and *M*_*b*_ localizations of particle *b* as:

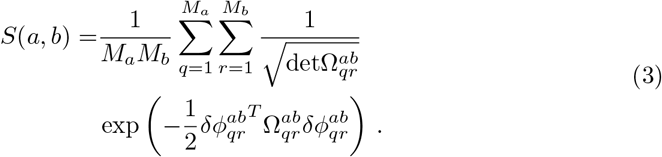

Here 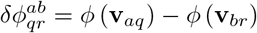 is the difference in transformed (rotated and translated) coordinates of localization *q* of particle *a* and localization *r* of particle *b*, and 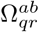 is defined in terms of the uncertainty covariance matrices of the localizations as:

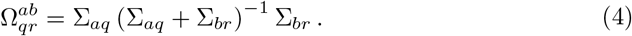

For example, for 2D-data with isotropic localization uncertainties, this reduces to:

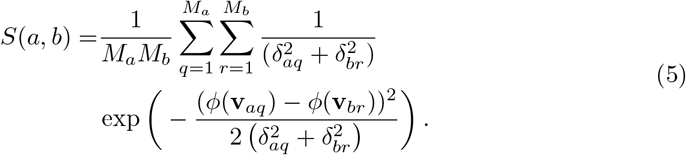

The normalization of the cost function with the numbers of localizations per particle reduces the impact of the variations in these number, which makes it a better descriptor of the similarity between the structure of the particles. The next step is to transfer dissimilarity values:

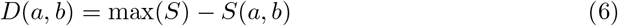

to spatial coordinates in a multidimensional space suitable for classification using multidimensional scaling (MDS) [19]. The transformed particles will then be partitioned into clusters by k-means clustering [3, 15] in this multidimensional space. Parameter settings for the classification step are given in section 3.2. This process is repeated for the JRMPC reconstructions *l* = 1, 2, …, *L* leading to *n* = 1, 2, …, *n*_*l*_ clusters that are denoted as 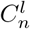(see Figure 1).

### 2.3 Connection

As we repeat the JRMPC reconstruction *L* times, pairs of clusters from different initializations may share different particles. Therefore, we need to combine the different clusters into a single well-aligned structure. In a first step we discard clusters with less than *ϑ* particles. This threshold helps to filter out poorly aligned clusters as well as clusters with particles of poor quality, as these tend to accumulate in clusters with low number of particles.

Next, the cluster with the largest number of particles is selected as initial estimate of the super-particle reconstruction **C**_*a*_. This main cluster, 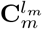, is used as the target for a pairwise comparison of clusters. A loop over all clusters 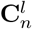 for *l* ≠ *l*_*m*_ is done, and clusters 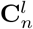 and 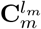 are compared to check for particles that are in both clusters. If there exists at least one common particle *c* with coordinates **V**_*c*_ ∈ **V** then the clusters 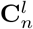 can be added to the super-particle reconstruction estimate **C**_*a*_ following:

Step 1: apply the inverse transformation of particle *c* in cluster 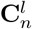 to transform all particles in the cluster 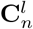 to the original position and pose of **V**_*c*_:

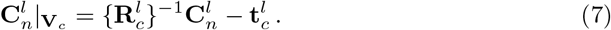

Step 2: apply the transformation of particle *c* in the main cluster 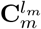 to all particles in the cluster 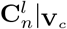 to the position and orientation of 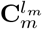:

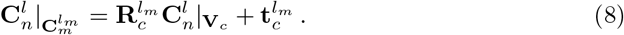

Now that the cluster 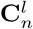 is aligned with the pose of the main cluster 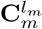 the particles of 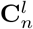 can be added to the super-particle reconstruction estimate **C**_*a*_. In this way more and more particles accumulate in the final reconstruction, yielding the final outcome of our proposed algorithm.

Care must be exercised for two subtleties. First, it can happen that there is more than one common particle between the two clusters 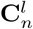 and 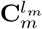. Then, if there exists more than one common particles between two clusters, we will calculate all the common particles’ translation matrices and rotation matrices from the cluster 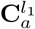 to the cluster 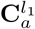,

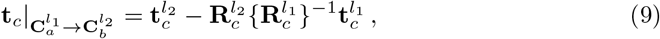

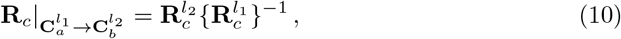

then we compare all the **t**_*c*_ and **R**_*c*_ and use the common particle with rotation and translation matrix that are closest to the median of all translation and rotation matrices of all the common particles. Second, we must check if the particles of cluster 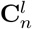 are not already in the reconstruction estimate **C**_*a*_. Only the unique particles that are not already contained in the reconstruction are added to **C**_*a*_.

The connection pipeline is summarized below:

#### Algorithm 1 Connection Algorithm

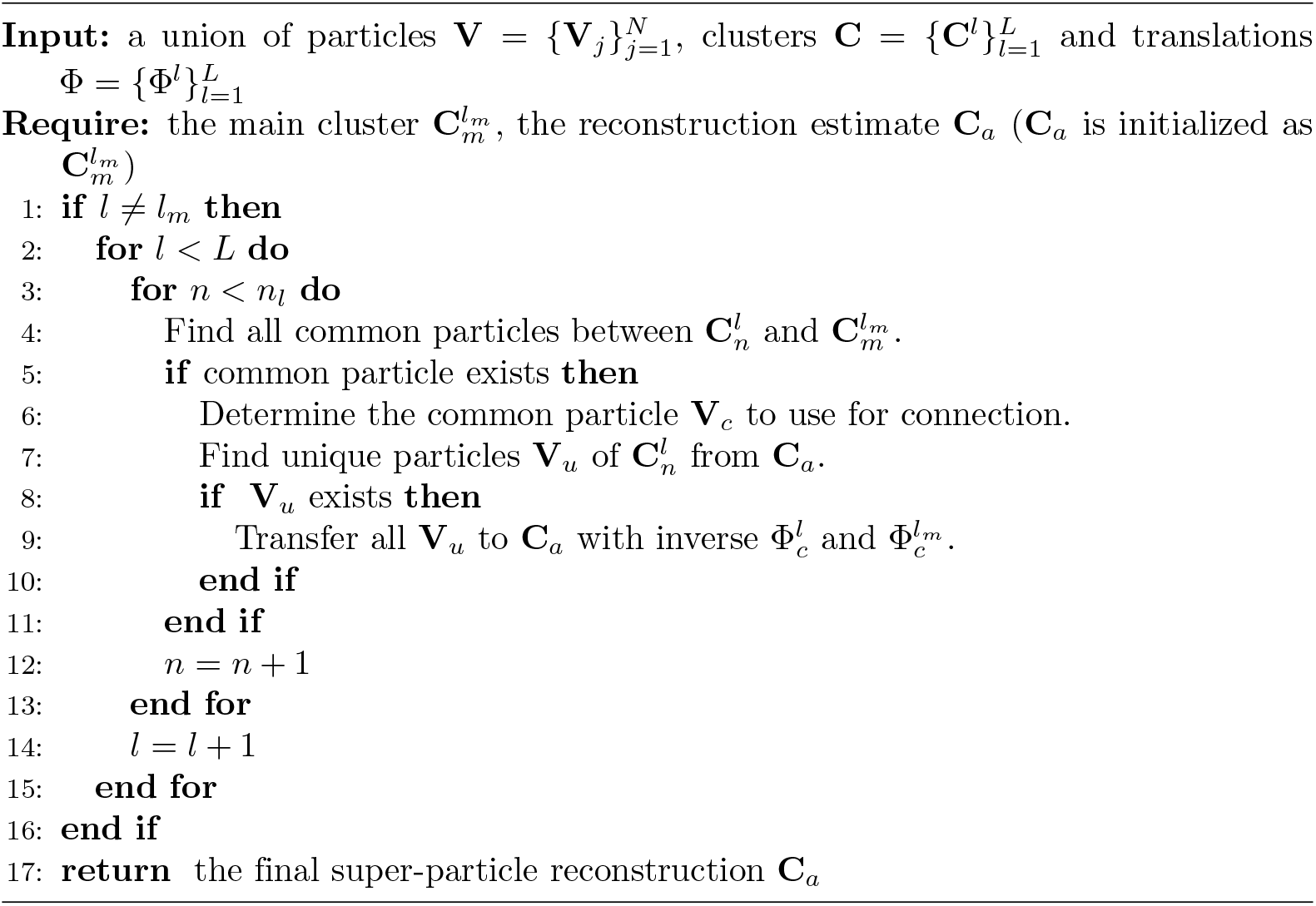

## 3 Experiments

### 3.1 Experimental Data

We applied our method to four different localization microscopy experiments and one simulation described here:

#### 3.1.1 DNA origami TUD-logo

We tested three different 2D TUD-logo DNA origami datasets [13] with *DOL* of 30%, 50% and 80%. We compared the results of the currently proposed method and the all-to-all method [13] in Figure 2 and Figure 5. The data is available online [12].

**Figure 2.**
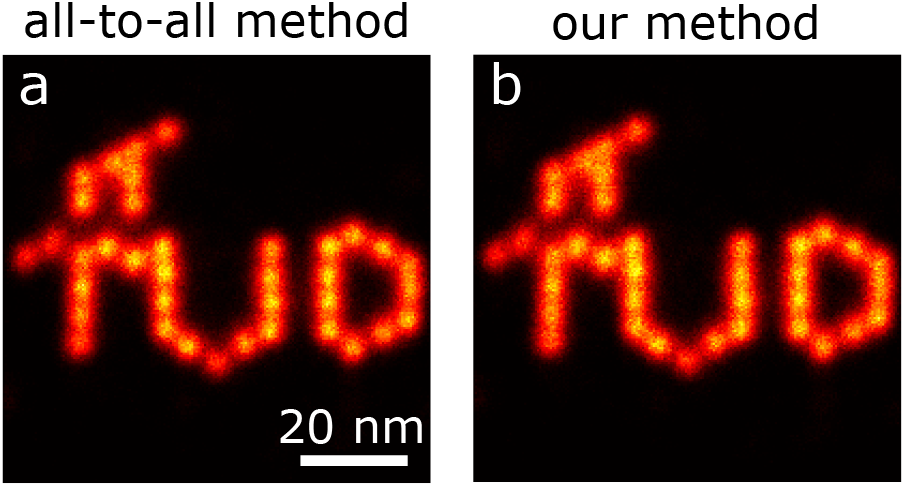
Comparison of particle fusion speed by our method with the all-to-all method on 383 experimental 2D TUD-logo DNA origami particles with *DOL*=80% and 788,875 localizations. (a) Reconstruction by all-to-all registration (FRC resolution 3.3 ± 0.3 nm, computational time about 2 hours (GPU)). (b) Reconstruction by our method (FRC resolution 3.2 ± 0.1 nm, computational time 9.5 minutes (CPU)).Scale bar applies to both images.

#### 3.1.2 2D nuclear pore complex

We further applied our method to 2D Nuclear Pore Complex (NPC) data which were previously described in ref. [18]. In Figure 4, we show our reconstruction of NPCs together with the reconstruction of the all-to-all method [13] to compare the methods’ capabilities in the reconstruction of symmetrical structures.

#### 3.1.3 3D nuclear pore complex

We applied our algorithm to 3D NUP107 NPC data [10] acquired by two different localization microscopy techniques. The data is available online [11]. The poses of the NPCs are experimentally constrained as they are all embedded in the nuclear envelope which is imaged as flat as possible on the cover glass. The lower and upper ring of all particles are therefore roughly perpendicular to the optical axis of the microscope [10].

#### 3.1.4 DNA origami Digits data

The so-called nanoTRON datasets [1] consist of DNA origami structures in the shape of the digits 1, 2, and 3 and in the shape of a 3 × 4 rectangular grid. The data is available online [9] and contains on the order of a few thousand particles. These datasets are used to showcase the processing speed advantages of our method.

#### 3.1.5 Simulation data

Simulation data of the DNA-origami TUD-logo was generated as described in [13].

### 3.2 Parameter Settings

A number of parameters in the three algorithmic steps of alignment, classification and connection must be set. The default values given in Table 1 are suitable for most of the cases.

**Table 1.** Parameter Settings

| definition | notation | default value |
| --- | --- | --- |
| # particles used to estimate $K$ | $\zeta$ | $\min(20, N)$ |
| # GMM centers used to estimate $K$ | $K_0$ | $\min(\frac{\sum_{j=1}^N M_j}{N}, 100)$ |
| # GMM centers | $K$ | depends on input |
| prior probability of $G_k$ | $p_k^0$ | $1/K$ |
| initial mean of $G_k$ | $\mu_k^0$ | randomly generated |
| initial rotation matrix | $\mathbf{R}_j^0$ | $\mathbf{I}_d$ |
| initial translation matrix | $\mathbf{t}_j^0$ | $\bar{\mu}^0 - \bar{\mathbf{v}}_j$ |
| initial standard deviation of $G_k$ | $\sigma_k^0$ | depends on input |
| # repetitions | $L$ | 2 |
| # clusters | $n_l$ | 2 |
| threshold for good cluster | $\vartheta$ | $N/(n_l + 1)$ |

We estimate the number of initial GMM centers *K* by applying the mean-shift method [3,6] to the outcome of *ζ* randomly selected input particles coarsely transformed by JRMPC with *K*_0_ randomly generated GMM centers. We set 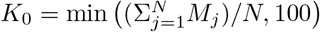, i.e. equal to the average number of localizations of all input particles with a minimum of 100. We choose *ζ* = 20, if the number of input particles *N* < 20 then *ζ* = *N*. The value of *K* estimated in this way is approximately equal to the number of binding sites in most cases. All initial values for the prior probabilities of the *K* Gaussians are set uniformly to *p*_*k*_ = 1*/K*. The initial values of the center positions 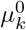 are generated randomly within a rectangular bounding box containing all the localizations. We initialize the transformation as 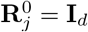 and 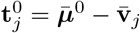, where 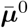 is the average of the *K* GMM centers. The diagonal of the bounding box containing all the input particles after applying the initial translation is set as the initial value of all Gaussian standard deviations 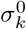. We set the default value for the number of clusters *n*_*l*_ to 2 in the classification step because the registration of JRMPC usually only contains two flipped structures. The threshold *ϑ* for a cluster to be used in the connection step is set as *N/*(*n*_*l*_ + 1). The default number of repetitions *L* for the JRMPC initializations is 2.

We use the default parameter settings throughout with two exceptions. The reconstruction of the nanoTRON 3 × 4 grid (Figure 3(d) uses non-default parameters with a larger number of clusters (*n*_*l*_ = 8) to guarantee clusters that contain well-aligned particles. The reconstruction of the 3D NPC particles (Figure 6) uses a non-default value for the initial Gaussian standard deviation (we use 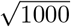, much smaller than the default value) to better fit with the limited range of initial poses of the NPCs. An inferior alignment is observed with the default value. In general we find that the quality of the individual clusters can be improved by increasing *n*_*l*_ or *ϑ*. A larger number of JRMPC initializations *L* can help to increase the number of particles in the final reconstruction after the connection step.

**Figure 3.**
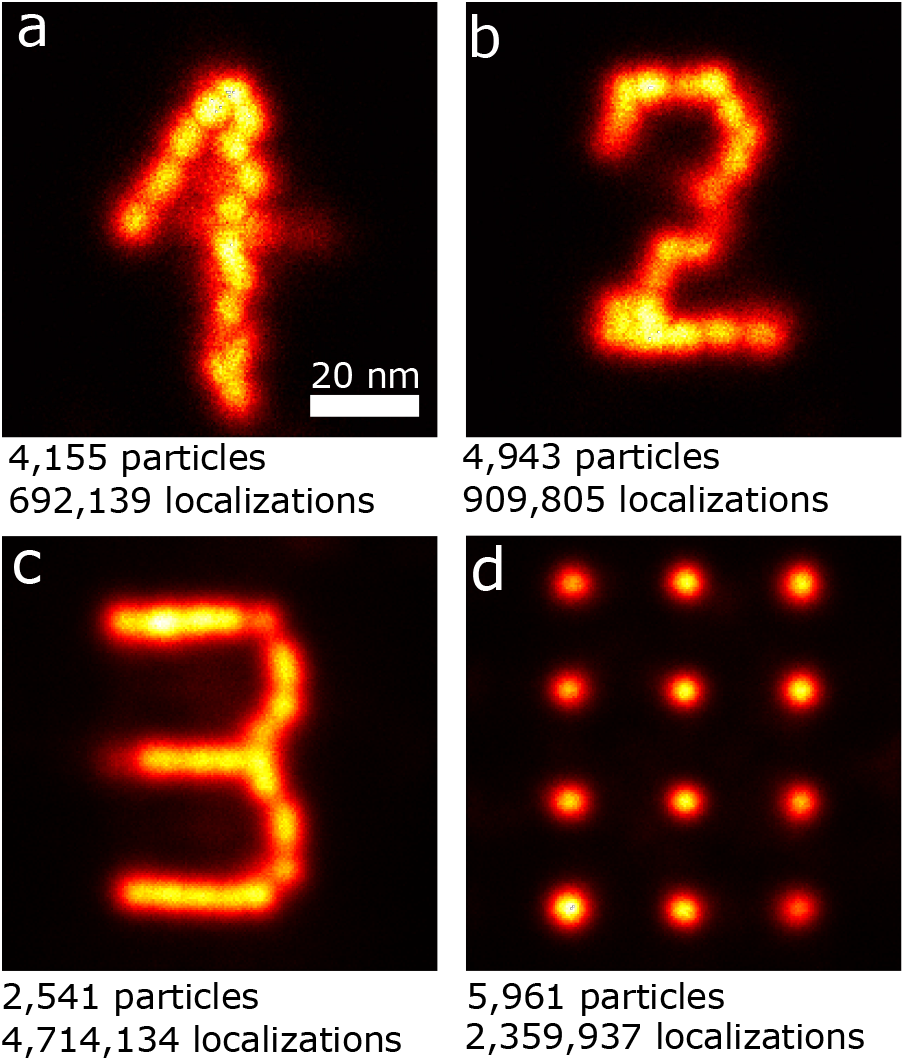
Particle fusion speed for experimental 2D DNA-origami with large amount of particles. (a) Reconstruction of digit 1, computational time 1.1 h (CPU). (b) Re-construction of digit 2, computational time 1.3 h (CPU). (c) Reconstruction of digit 3, computational time 48 min (CPU). (d) Reconstruction of 3 × 4 grid, computational time 4.8 h (CPU). The number of particles and localizations in each reconstruction are indicated below the figures. Scale bar of (a) applies all sub-images.

### 3.3 Benchmark Algorithms and Evaluation Metrics

We compare our proposed method with the all-to-all method [13] [10]. We use the Fourier Ring Correlation (FRC) [21] to measure the resolution of the super-particle reconstructions. We form two independent input image subsets from the super-particle reconstruction to perform the FRC analysis. The first subset is the main cluster 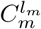 and the second subset consists of all other particles in the reconstruction. These two subsets can be used as statistically independent image subsets that are the necessary inputs for the FRC measurement because each subset contains a similar number of different particles from different independent experiments. We cross-checked the outcomes of this FRC computation with the standard method of independently processing two subsets of the total set of input particles and found outcomes within the uncertainty margin of the FRC estimation. In addition, we calculate the localization distribution over the azimuthal angles to analyze the reconstruction symmetry for symmetrical structures. For the 3D NPC data, we also visualize and compare the distributon of *z* positions of the localizations, the radius of each of the two rings, and in a rose plot the localization distribution over azimuthal angles. In the simulations, we compute the root mean square distance between the localizations after particle fusion and the attendant binding sites to quantify the quality of the fusion process [10].

## 4 Results

### 4.1 Computational Cost

Compared to the all-to-all method, which has an unfavorable computational cost scaling as *N*^2^, our method is much faster as it is linear with *N*. Figure 2 shows the reconstructions of 383 experimental TUD-logo particles with *DOL*=80% and 788,875 localizations obtained with the all-to-all method and our method. We repeated our method on the 80% *DOL* TUD-logo particles 30 times in order to assess the uncertainty in FRC-resolution and computation time. Both methods achieve a similar reconstruction quality, consistent with near equal FRC resolutions (3.3 ± 0.3 nm for the single instance of the all-to-all, 3.6 ± 0.3 nm for the 30 runs for our method). The computational time of the all-to-all method, however, is almost 12 times longer than for our method. More importantly, our computational time of 9.6 ± 0.6 minutes was performed on a simple CPU (40 core Xeon E5-2670v3), opposed to the GPU-implementation of the all-to-all registration (K40c Tesla GPU). The all-to-all method is practically impossible on a CPU when having more than 100 particles. The estimated number of Gaussian centers *K* is 40±3, which is close to the actual number of binding sites (37). The random initializations of the JRMPC usually result in a final GMM that is similar to the combination of two inverted TUD-logos, which can be classified appropriately in only two clusters. Our method can effectively handle large amounts of particles because of the favorable reconstruction speed. To show the capability of our method to handle this large data we applied it to the nanoTRON datasets, which contain an order of magnitude more particles than the TUD-logo datasets. We achieved clear structures of the digits 1, 2, and 3 and of the 3 × 4 grid in only 1.1 h, 1.3 h, 45 min. and 4.8 h, respectively, in CPU compared to a computational time of multiple days for the GPU-accelerated all-to-all method. It would have taken several days to resolve the full dataset with the all-to-all method. Due to this speed limitation we only used part of the data in the all-to-all method. The FRC resolution obtained by the all-to-all registration for these four datasets (digits 1, 2, and 3 and of the 3 × 4 grid) containing 1219, 1309, 1278 and 1194 particles are 3.69 ± 0.02 nm, 4.40 ± 0.19 nm, 3.98 ± 0.22 nm and 3.59 ± 0.15 nm, respectively [14]. Our reconstructions include 4155, 4943, 2541 and 5961 particles for these four datasets and the FRC resolutions are 2.76 ± 0.92 nm, 2.80 ± 0.54 nm, 3.21 ± 0.33 nm and 3.51 ± 0.28 nm, respectively. These numbers are smaller as we are able to assemble more particles in the final reconstruction compared to the all-to-all method. For the digits 1, 2, and 3, the estimated *K* (25, 23, 34) is close to the actual number of binding sites (18, 23, 25). For the 3 × 4 grid particles, our *K*-estimation algorithm estimates *K* = 42 which is much more than the 12 binding sites. For that reason the JRMPC reconstructions have more clusters and we need a larger *n*_*l*_ = 8 to separate them correctly.

### 4.2 2D NPC data: influence of symmetry

Our method also overcomes the second disadvantage of the all-to-all method, the hot-spot problem occurring for symmetrical structures. In Figure 4, we compare reconstructions of 2D NPC particles with eight-fold rotational symmetry. The reconstruction of the all-to-all method without prior knowledge (Figure 4(a)) and (b)) shows one apparent “hot-spot” with more than 600 localizations compared to other blobs with around 400 localizations. After imposing eight-fold rotational symmetry the hot-spot disappears (Figure 4(c)). Imposing this symmetry changes the ellipticity of the reconstructed NPC ring from the earlier 0.89 to 0.99. So, symmetry has been restored, but at the expense of a shape that changed from an ellipse to a circle. Our method × applied to the same NPC particles does not result in a hot-spot (Figure 4(e)), quantified by a more uniform distribution of localizations over the 8 peaks (compare (b) and (f)). The ellipticity of our reconstruction is 0.86 which matches reasonably well with the all-to-all value of 0.89. The number of Gaussian components *K* in the GMM is estimated by our algorithm to be 8 which is obviously equal to the number of visible binding sites in the 2D NPC.

**Figure 4.**
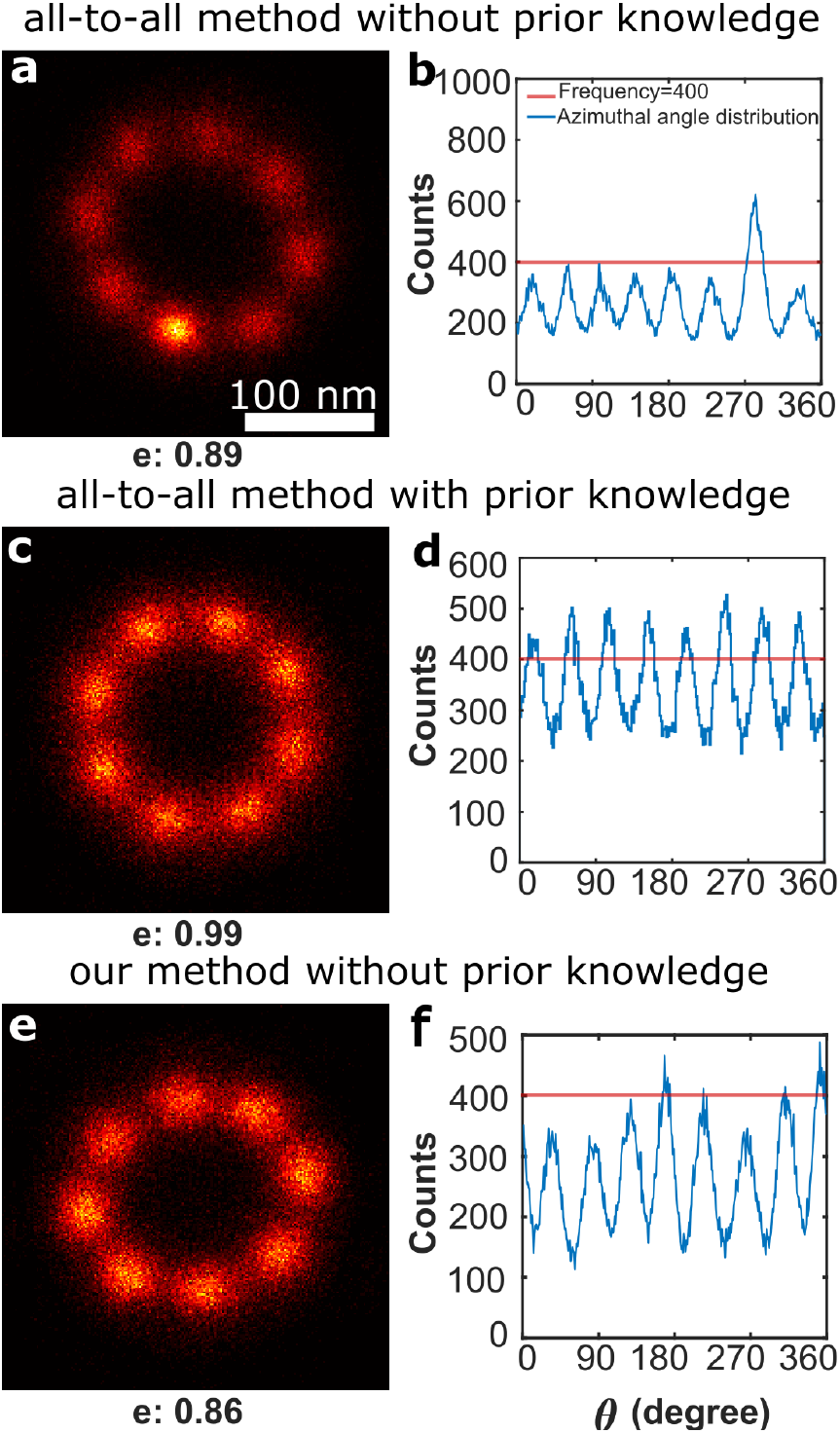
Comparison of particle fusion performance between our method and the all-to-all method on 304 experimental 2D nuclear pore complex particles. (a) Reconstruction with the all-to-all method without prior knowledge. A “hot-spot” is visible due the enhancement by pair-wise registration. Fitted ellipticities *e* to the reconstruction are shown below.(b,d,f) Histogram of the azimuthal angles of the localizations in (a,c,e) respectively; for comparison, a red line indicates 400 counts. (c) Reconstruction with the all-to-all method after explicitly imposing eight-fold symmetry. (e) Reconstruction with our method without prior knowledge. Even without imposing symmetry no hot-spot occurs. Scale bar of (a) applies to (c,e).

### 4.3 Low labelling 2D DNA origami data

A major accomplishment of the all-to-all method is its ability to handle poorly labelled data. It appears our method outperforms the all-to-all method even in this respect. Figure 5 shows a comparison of reconstructions of hundreds of TUD-logos with low *DOL* values equal to 50% and 30%. Our method results in a visually better reconstruction quality, especially for the worst quality *DOL*=30% dataset (compare Figure 5(a) and (c)). Nearly all binding sites on the origami at a distance of about 5 nm are resolved in (c) where in (a) especially the edges are washed out and localizations are concentrated to a few binding sites. This is consistent with the FRC resolutions of 3.1 nm and 3.3 nm for the 50% and 30% *DOL* datasets, respectively, which compares favourably with the FRC resolutions for the all-to-all method equal to 3.5 nm and 5.0 nm for the 50% and 30% *DOL* datasets, respectively. The mean-shift method estimates *K* = 46 for the data with 30% *DOL* and *K* = 37 for 50% *DOL*. These two *K* values are very close to the actual number of 37 binding sites of the origami design. The initial Gaussian standard deviation is quite large (∼ 100 nm) at first. Most of the Gaussian components shrink to a small size (less than 3 nm) eventually, and only a few to a medium size (∼10 nm). Most of the initially randomly generated GMM centers ***µ***_***k***_ are finally positioned near the binding sites of the TUD-logo.

**Figure 5.**
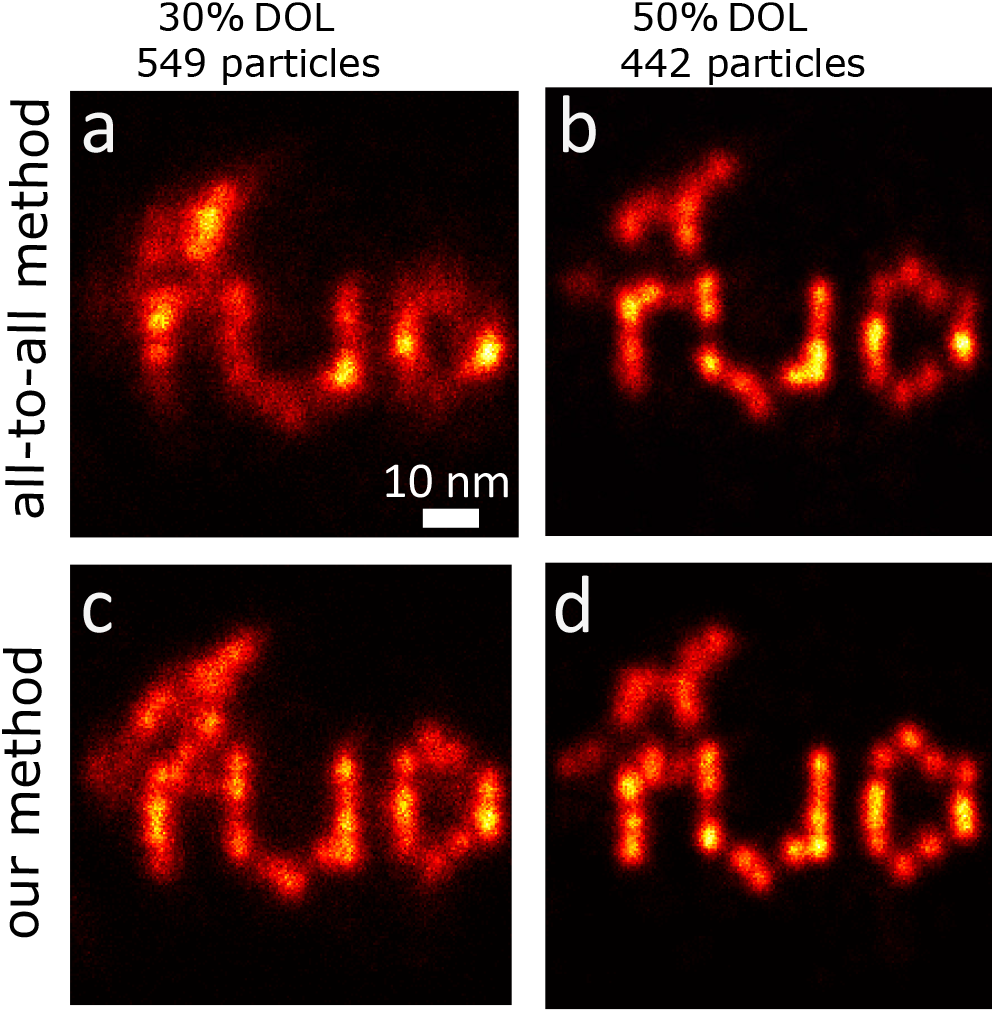
Comparison of the particle fusion performance with our method the all-to-all method on experimental 2D TUD-logo DNA origami particles with low density of labelling (*DOL*).(a-b) Reconstructions using all-to-all registration (FRC resolution of 5.0, 3.5 nm for 30% and 50% *DOL* respectively). (c-d) Reconstructions using our method (FRC resolution of 3.3, 3.1 nm for 30% and 50% *DOL* respectively). Scale bar of (a) applies to (b-d).

### 4.4 3D NPC data

Another major achievement of the all-to-all method is the ability to reconstruct 3D data [10]. Our method shows a comparably good performance on 3D datasets. Figure 6 shows a comparison of 3D Nup107 NPC structures imaged with both PAINT and STORM. Our method shows reconstructions of similar quality as the all-to-all method (compare Figure 6(a) and (k) and compare Figure 6(f) and (p)). Here, the all-to-all method relies on detecting the rotational symmetry from the data and subsequently promoting the symmetry in the reconstruction. In contrast, neither prior knowledge or detection of symmetry nor extra post-processing is needed with our method. Comparison of Figure 6(b,g,l,q), (c,h,m,l) to (d,i,n,s), respectively, further shows that our method obtains similar NPC structural parameters (the distance between the nuclear and cytoplasmic rings and their radius) as the all-to-all method. The rose plots Figure 6(e,j) obtained from the all-to-all method’s reconstructions show eight-fold symmetry for each ring, and the number of localizations in each peak is almost the same. The rose plots Figure 6(o,t) of our reconstructions also clearly show eight peaks for each ring, but the number of localizations in each peak is slightly different. This is reasonable considering that our method does not rely on symmetry in the reconstruction. Our *K*-estimation algorithm estimates *K* = 34 for both cases, which is also reasonable as the number of actual binding sites should be 32 given the structure of the EM model [17, 24]. The default value of *σ*_*k*_ does not work here and we used 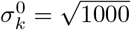 nm instead. The final center points of the GMMs are nearly all distributed inside the 16 spheres of the 3D NUP reconstructions.

**Figure 6.**
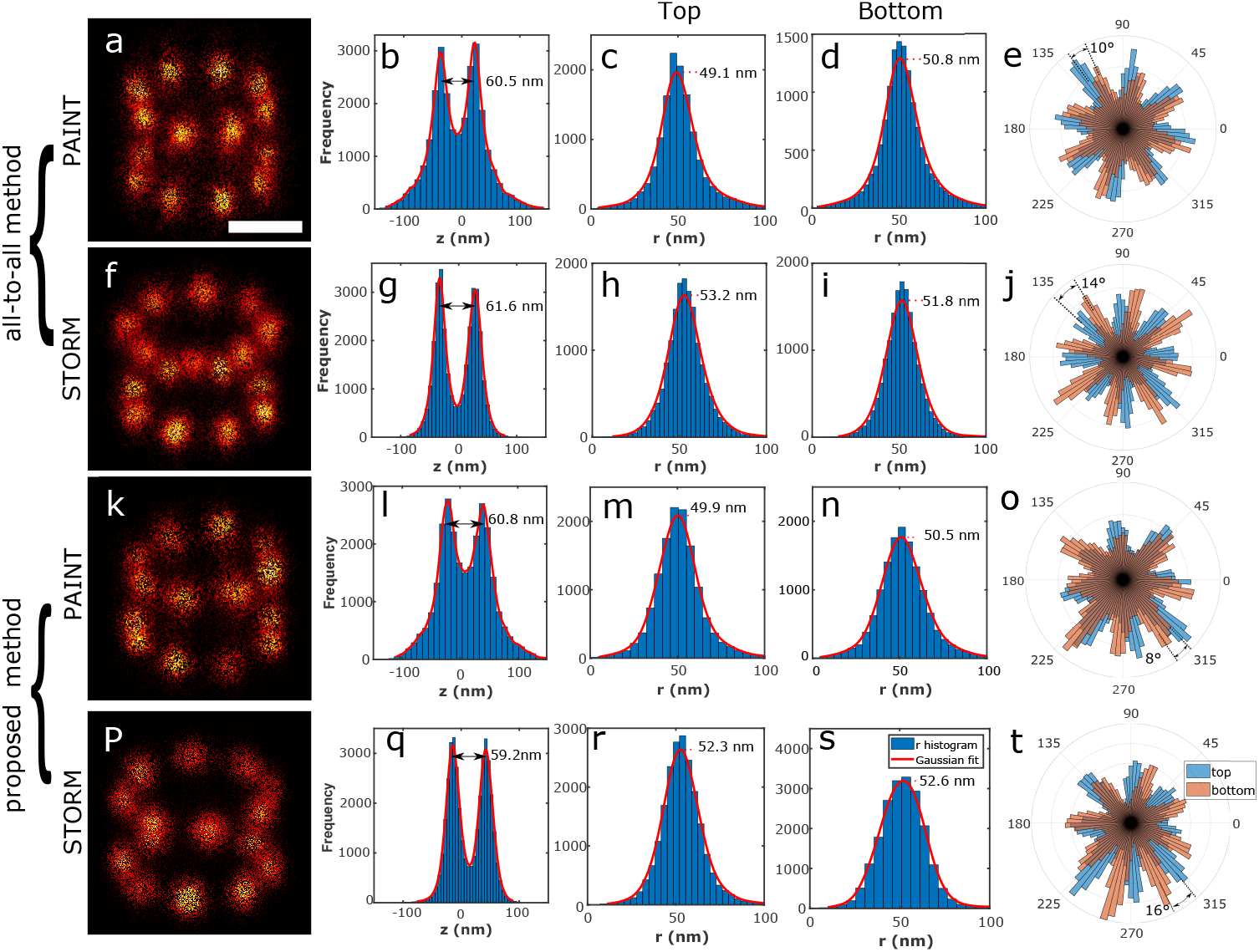
Comparison of particle fusion performance between our method and all-to-all method on experimental 3D Nup107 particles acquired by different localization microscopy techniques. (a) Fusion of 306 Nup107 particles obtained from 3D astigmatic PAINT reconstructed by the 3D all-to-all method. (b,g,l,q) Histogram of the *z* coordinate of localizations in the reconstruction (a). (c,h,m,r) Histogram of the radius of upper ring’s localizations, (d,i,n,s) lower ring. (e,j,o,t) Rose plot of the localization distribution over azimuthal angles for the upper and lower rings of the reconstructions. (f) Fusion of 356 Nup107 particles obtained from 3D astigmatic STORM reconstructed by the 3D all-to-all method. (k) Fusion of 306 Nup107 particles obtained from 3D astigmatic PAINT reconstructed by our method. (p) Fusion of 356 Nup107 particles obtained from 3D astigmatic STORM reconstructed by or method. Scale bar indicates 50 nm and applies to a,f,k and p.

## 5 discussion

We explore the limitations of the proposed method in terms of *DOL*, localization precision and the number of particles by applying our method on simulated TUD-logo datasets. These simulated data have the default settings of 200 particles, 2000 detected photons per localization event (corresponding to an average localization uncertainty of 4.85 nm) and 60% *DOL*. When we change one of these three parameters, we keep the other two at the default values. We simulate more challenging conditions than most often encountered in real experiments to probe the performance limitations. We perform ten independent simulations for each setting of the simulated data. We evaluate the reconstruction quality by calculating the average distance of localizations to the corresponding binding sites (*AD* in short) following ref. [10] where this measure was introduced for simulated data. For a simulated structure with around 5 nm distance between binding sites corresponding to an DNA Origami design, then an error of *AD* < 10 nm is needed for the reconstruction to appear reasonably correct; for *AD* < 5 nm, binding sites details can be observed in the reconstruction. Figure 7(a) shows that the error (*AD*) decreases with increasing *DOL*. For *DOL* larger than about 40% our method can stably obtain a clear reconstruction.

**Figure 7.**
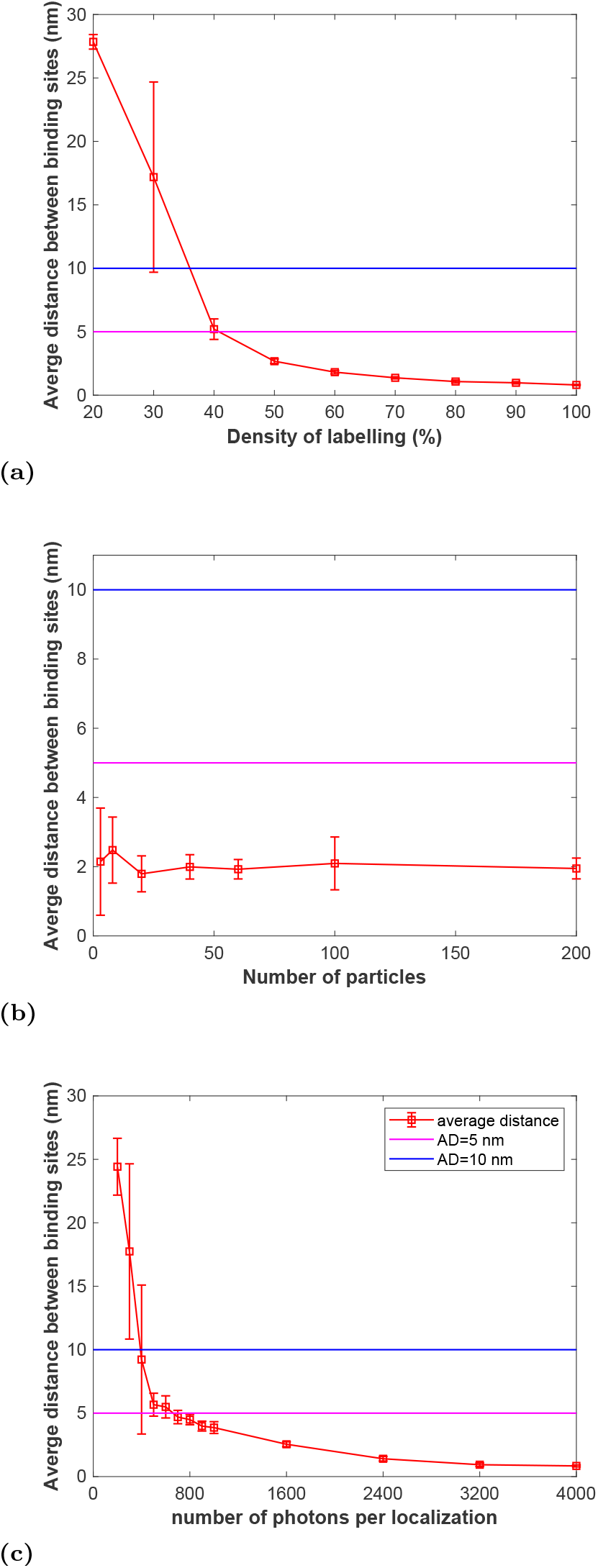
Simulation study of limitation of the proposed method. Each point in the graph indicates ten independent experiments. Reconstructions with *AD* < 10 nm (magenta line) are assessed as ‘correct’ and with *AD* < 6 nm (blue line) as ‘clear’. (a) Reconstruction quality as a function of *DOL* for 200 particles with 2000 photons per localization. (b) Reconstruction quality as a function of number of input particles with 60% *DOL* and 2000 photons per localization. (c) Reconstruction quality as a function of number of photons per localization for 200 particles with 60% *DOL*.

Our method is not sensitive to the number of input particles. For particle numbers varying from 3 to 200, *AD* values are always less than 5 nm and fluctuate in a small range (Figure7(b)). Even though the registration for input particles less than 10 appears correct, the underlying structure is still hardly visible in the reconstruction because of the small total number of localizations.

Figure 7(c) indicates that the error *AD* decreases with increasing number of photons per localization. With 200 input particles with 60% *DOL*, our method is able to Rose plots in (e,j) show 8 fold symmetry with nearly equal number of localizations, but symmetry was used here in the reconstruction. Rose plots (o,t) Without any prior knowledge reconstruction with our method also shows 8 clear peaks however with a stronger variation in the number of localizations. correctly reconstruct the underlying structure as long as the number of photons is greater than 400, corresponding to a localization uncertainty of 12 nm which is 2.4 times larger than the minimum binding site distance 5 nm.

Several of the results we obtained can be qualitatively understood:

In comparison to the all-to-all-method our approach produces better results for poor, underlabeled data. The reason is that in the pairwise registration of the all-to-all method pairs of poor quality particles must be aligned, which is more error prone than our approach where each of the poor quality particles is aligned to the average of all particles. The same line of reasoning applies to the case of symmetric structures. The pairwise registration of the all-to-all method aligns random peaks that occur through the stochastic variations of labeling within the particles, while for our approach each particle is aligned to the average of all particles which smoothens out the stochastic variations in labeling.

We attribute the JRMPC local optima that consist of several distinct clusters with different poses to a difference in convergence rate between the widths of the Gaussian components and the particle rotations. It seems that the Gaussian widths shrink relatively fast, while the particle rotations only change slowly, as the iteration progresses. This results in posterior probabilities *α*_*kij*_ for the Gaussian component *k* that is nearest to localization *i* of particle *j* that quickly converge to nearly one and to virtually zero for the other Gaussian components. On the other hand, for the case of 3D NPC particles with a limited range of poses in the dataset, the widths of the Gaussian components appear too large, leading to sets of particle rotations that are distributed too broadly. Summarizing, the reconstruction quality appears to be sensitive to the initial setting and convergence rate of the Gaussian widths.

A number of algorithmic improvements can be envisioned. First of all we could incorporate the localization uncertainties in the JRMPC method, such that the probability of localization *i* of particle *j* to fit Gaussian component *k* is a normal distribution with a variance that is the sum of the variance due to the localization uncertainty and the variance of the Gaussian component. Especially in cases where the localization uncertainty is on the order of the distance between binding sites, or where there is a broad distribution of localization uncertainties, or when the localization uncertainty is anisotropic (for 3D datasets), this may improve the sensitivity to the initial setting of the widths of the Gaussian components, as well as promote convergence to a global optimum. Another improvement may be found in a better description of the quality of the clusters. Now we opt for the simple criterion of number of particles in the cluster. Using the FRC resolution may be a better practice for assessing cluster quality.

## 6 Conclusion

We have proposed a fast particle fusion method with computational complexity that scales linearly with the number of input particles. In our method we apply the JRMPC method for multiple initializations and then use classification and connection steps to generate a correct reconstruction with as many particles as possible. The reconstruction quality of our method is measured by the FRC resolution and compared with the all-to-all method, revealing that our results are of comparable or better quality. Our method is fast, even without GPU acceleration, avoids symmetry artifacts, applies to 2D and 3D datasets, and reconstructs poor data with a limited number of particles, a low density of labelling and a large localization uncertainty.

## Acknowledgements

We thank Sabri Bolkar for initial attempts to apply the JRMPC method to single molecule localization microscopy.

## Appendices

### A Summary of JRMPC

The JRMPC method [4] is cast as an Expectation Maximization (EM) algorithm. In this framework the observed data are the set of particles *j* = 1, 2, …, *N* with localizations *i* = 1, 2, …, *M*_*j*_ represented by coordinates **v**_*ji*_. The localization uncertainties are not taken into account in the JRMPC method. The estimated parameters are the parameters of the *K* Gaussians of the GMM, and the rotations and translations of the particles that best match the GMM, defined as Θ in Equation 1. The latent or unobserved data *Ƶ* concern the assignment of localizations *i* in particle *j* to Gaussian *k* of the GMM. We have modified the original approach of ref. [4] by ignoring the outlier probabilities, i.e. there is no outlier class where the localizations can be assigned to. The expectation value of the log-likelihood over the distribution of latent data can be expressed as the sum over Gaussians *k*, particles *j* and localizations *i* as:

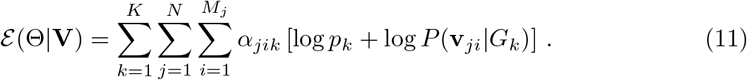

Here, the marginal probability that localization *i* of particle *j* fits Gaussian *k* is given by the normal distribution:

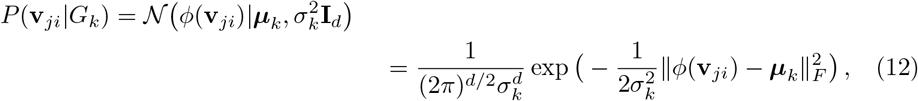

where ‖ · ‖ _*F*_ denotes the Frobenius norm, the Gaussian weight *p*_*k*_ is the probability *p*_*k*_ = *P* (*G*_*k*_ Θ), and the coefficients *α*_*jik*_ represent the posterior probability of the latent variable, i.e. the probability that localization *i* of particle *j* is assigned to Gaussian *k*. Starting point of each iteration round of the JRMPC is to update the posterior probabilities according to:

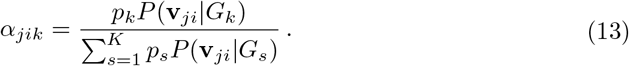

The next step is the update of the rotation and translation matrices. It appears that finding the optimum log-likelihood expectation value can be cast as:

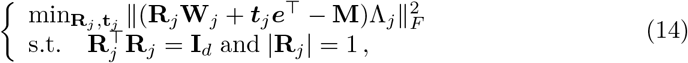

where **Λ**_*j*_ ∈ ℝ^*K×K*^ is a diagonal matrix with elements 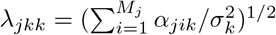, ***e***∈ ℝ^*K*^ is a vector of ones, **M** = [***µ***_1_, …, ***µ***_*K*_] ∈ ℝ^*d×K*^ represents a matrix of the means of the Gaussian components, and **W**_*j*_ = [***w***_*j*1_, …, ***w***_*jK*_] ∈ ℝ^*d×K*^ is the weighted-average localizations of the *j*^*th*^ particle, where ***w***_*jk*_ represents the single weighted-average localization of the *j*^*th*^ particle assigned to the *k*^*th*^ Gaussian component

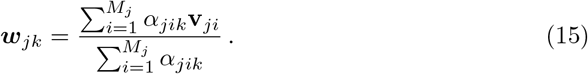

The optimal transformations 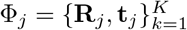 are subsequently found using the method of Umeyama [25]. Then, the optimal means and covariances of the Gaussian components are estimated. It turns out that the closed-form expressions:

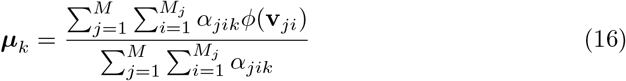

and

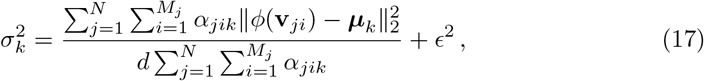

provide the sought-for optimum. Here, a small scalar *ϵ* is added to avoid singularities of *λ*_*jkk*_, as could arise if a Gaussian component has near zero localizations with a substantial posterior probability *α*_*jik*_.

Finally, the weights of the Gaussian components are updated according to:

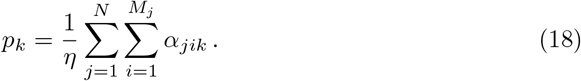

The different steps of the iterative JRMPC procedure are summarized as below:

#### Algorithm 2 JRMPC Algorithm

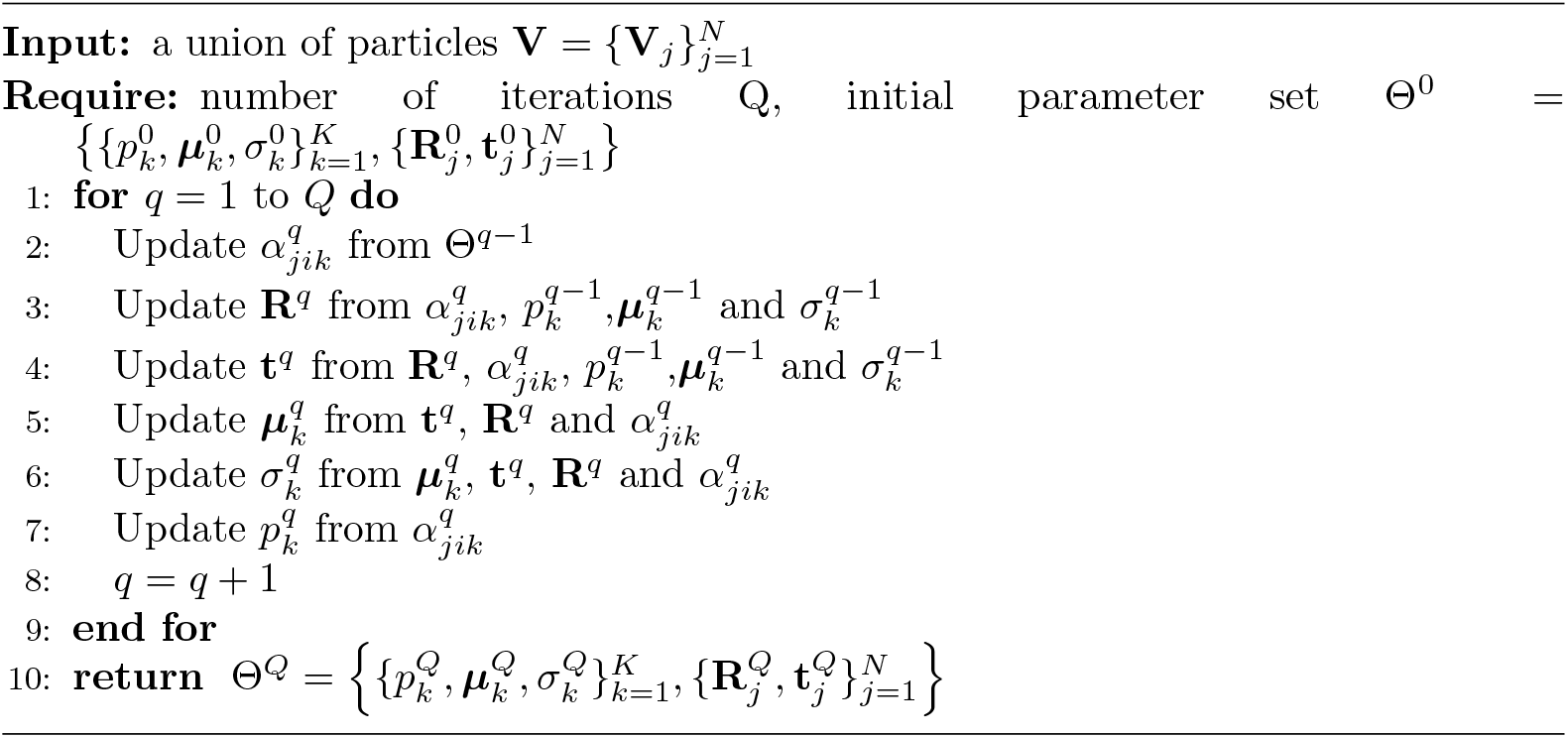

